# Knock-out of *CmNAC-NOR* affects melon climacteric fruit ripening

**DOI:** 10.1101/2022.02.02.478821

**Authors:** Liu Bin, Miguel Santo Domingo, Carlos Mayobre, Ana Montserrat Martín-Hernández, Marta Pujol, Jordi Garcia-Mas

**Author notes:** Corresponding authors: Jordi Garcia-Mas, Marta Pujol.

## Abstract

Fruit ripening is an important process that affects fruit quality. In melon, *ETHQV6*.*3*, a QTL involved in climacteric ripening regulation, was previously found to be encoded by *CmNAC-NOR*, a homologue of the tomato *NOR* gene. To further investigate *CmNAC-NOR* function we have obtained two CRISPR/Cas9 mediated mutants (*nor-3* and *nor-1*) in the climacteric Védrantais background. *nor-3*, containing a 3-bp deletion altering the NAC domain A, resulted in ~8 days delay of ripening without affecting fruit quality. In contrast, *nor-1* contained a 1-bp deletion resulting in a fully disrupted NAC domain, which completely blocked climacteric ripening. *nor-1* fruits did not produce ethylene, abscission layer was not formed and there was no external color change. Additionally, volatile components were dramatically altered, seeds were not well developed and flesh firmness was also altered. *nor-1* allele in heterozygosis showed ~20 days delay of fruit ripening. Our results provide new information regarding the function of *CmNAC-NOR* in melon fruit ripening, suggesting that it as a potential target to modulate shelf life in climacteric melon commercial varieties.

## Introduction

Fruit maturation is an important developmental stage because the ripening process activates a set of biochemical pathways that make the fruit attractive, perfumed, and edible (Bouzayen et al., 2010). In addition, the ripening process also helps plants spread the seeds (Wang et al., 2020a). Therefore, many efforts are ongoing to help understanding the complex regulation of this important trait (Bouzayen et al., 2010). Based on their ripening behavior, fleshy fruits have been divided into two groups: climacteric and non-climacteric (McMurchie et al., 1972). Climacteric fruits such as tomato are characterized by an ethylene burst accompanied by an increase in respiration at the onset of ripening. In contrast, non-climacteric fruits such as orange are characterized by a lack of the ethylene-associated respiratory peak (Paul et al., 2012). Usually climacteric fruits show shorter shelf-life than non-climacteric fruits (Hiwasa-Tanase and Ezura, 2014), and breeding programs in fruit crops are oriented to increase shelf life to minimize postharvest losses (Payasi and Sanwal, 2010).

Ethylene plays a primary role in initiating climacteric fruit ripening (McMurchie et al., 1972). In climacteric fruit, low ethylene production is present at the pre-climacteric stage, while an auto-stimulated massive ethylene production is present at the onset of the ripening stage. Exogenous ethylene treatment can also induce the ethylene burst at the pre-climacteric stage of climacteric fruits, thereby increasing the ripening process (Hiwasa-Tanase and Ezura, 2014). In contrast, treatment with the ethylene inhibitor 1-Methylcyclopropene (1-MCP) delays fruit ripening (Blankenship and Dole, 2003; Watkins, 2006). Antisense-induced repression of ethylene synthesis enzymes in tomato also delays fruit maturation (Hamilton et al., 1990). These results confirmed the key role of ethylene in regulating ripening in climacteric fruits. Interestingly, some reports showed that ethylene can also play a role in non-climacteric fruit ripening (Bouzayen et al., 2010; Katz et al., 2004), although in non-climacteric fruit a low level of ethylene during the whole developmental process is found (Paul et al., 2012). In melon some fruit ripening processes are independent of ethylene, as flesh softening, sugar accumulation or flesh color that did not change in ethylene-suppressed melon fruit (Flores et al., 2001; Pech et al., 2008), confirming that the control of fruit ripening is a complex trait.

Genes involved in fruit ripening have been largely studied in either climacteric or non-climacteric species (Gapper et al., 2013; Lü et al., 2018; Osorio et al., 2013) and tomato has emerged as a prime model of climacteric fruit ripening (Alexander and Grierson, 2002). Genetic characterization of several ripening related mutants in tomato has advanced our knowledge of the mechanisms that regulate fruit ripening (Giovannoni, 2007). The *ripening-inhibitor* (*rin*), *non-ripening* (*nor*) and *Colourless non-ripening* (*Cnr*) mutations have been useful in understanding the transcriptional regulation of fruit ripening (Giovannoni, 2007; Manning et al., 2006; Vrebalov et al., 2002; Wang et al., 2020b). In addition, a number of additional ripening related genes have been identified through reverse genetic approaches or transcriptomic studies (Gapper et al., 2013; Lin et al., 2008; Vrebalov et al., 2009). For non-climacteric fruit, one of the most studied plants has been strawberry (Osorio et al., 2013) and recent studies identified several genes that were involved in strawberry fruit ripening, including *FaPYR1* (Chai et al., 2011), *FaExp2* (Civello et al., 1999), *FaASR* (Chen et al., 2011), *FaABI1*(Jia et al., 2013), and *FaRIF* (Martín -Pizarro et al., 2021). These studies have provided valuable information on the gene function related to fruit ripening regulation.

Melon (*Cucumis melo* L.) is a suitable model to study fruit ripening, because it contains both climacteric and non-climacteric genotypes (Ezura and Owino, 2008). Genetic analysis on a biparental population of the cantaloupe type Védrantais (VED, climacteric) × PI 161375 (SC, non-climacteric) inbred lines indicated that ethylene production and fruit abscission were controlled by two independent loci, *Al-3* and *Al-4* (Périn et al., 2002). In recent studies, a near-isogenic line SC3-5-1 derived from the non-climacteric parental lines SC and the inodorus type Piel de Sapo (PS) showed a climacteric ripening phenotype, and two QTLs, *ETHQB3*.*5* and *ETHQV6*.*3*, were found to be involved in the regulation of climacteric ripening in SC3-5-1 (Eduardo et al., 2005; Moreno et al., 2008; Vegas et al., 2013). In a previous work, *ETHQV6*.*3* was found to be encoded by a *NAC* transcription factor *CmNAC-NOR*, phylogenetically related to the tomato *SlNAC-NOR* (Ríos et al., 2017). TILLING mutants containing non-synonymous mutations in the coding region of *CmNAC-NOR* showed a delayed ripening phenotype, suggesting that *CmNAC-NOR* is an important regulator of climacteric ripening in melon. To further investigate the *CmNAC-NOR* function, in this study we have generated and phenotyped CRISPR/Cas9 mutants with different disruption levels.

## Results

### Generation of *CmNAC-NOR* disrupted mutant lines by CRISPR/Cas9

*CmNAC-NOR* contains three exons and encodes a protein of 353 amino acids (Ríos et al., 2017). In this study, three guide RNAs (gRNAs) that specifically target the first or second exon of *CmNAC-NOR* were designed (Figure 1A). After melon transformation, we obtained 83 and 6 T_0_ plants containing the gRNA2-gRNA1-CAS9 and the gRNA1-gRNA2-CAS9 construct, respectively, but none of them were edited. In contrast, we obtained 39 T_0_ lines that contained the gRNA3-gRNA1-CAS9 construct, and six mutations at the gRNA3 target site were detected (Figure 1B). As melon tissue culture often induces the generation of tetraploid plants (Ezura et al., 1992), we looked at the ploidy of 15 individuals and found three diploid plants (Figure 1C). These diploid plants contained two different mutations, a -3 bp and a -1 bp deletion, which were named *nor-3* and *nor-1*, respectively (Figure 1D). The -3 bp deletion in *nor-3* results in the loss of the Proline in position 17 (Figure 1E), which is predicted as a deleterious change (score:-14.874) by PROVEAN (Protein Variation Effect Analyzer) (Choi and Chan, 2015). The -1 bp deletion in *nor-1* results in major changes from amino acid 16^th^ and the generation of a truncated protein of 37 aa (Figure 1E). The functional regions located in the NAC subdomain A of the CmNAC-NOR protein were totally disrupted in *nor-1* (Figure 1E), suggesting that *nor-1* is a loss-of-function mutant. In the following studies, we used T_1_ plants of the two lines. For the controls, we used VED and T_1_ non-edited (NE) plants.

**Figure 1.**
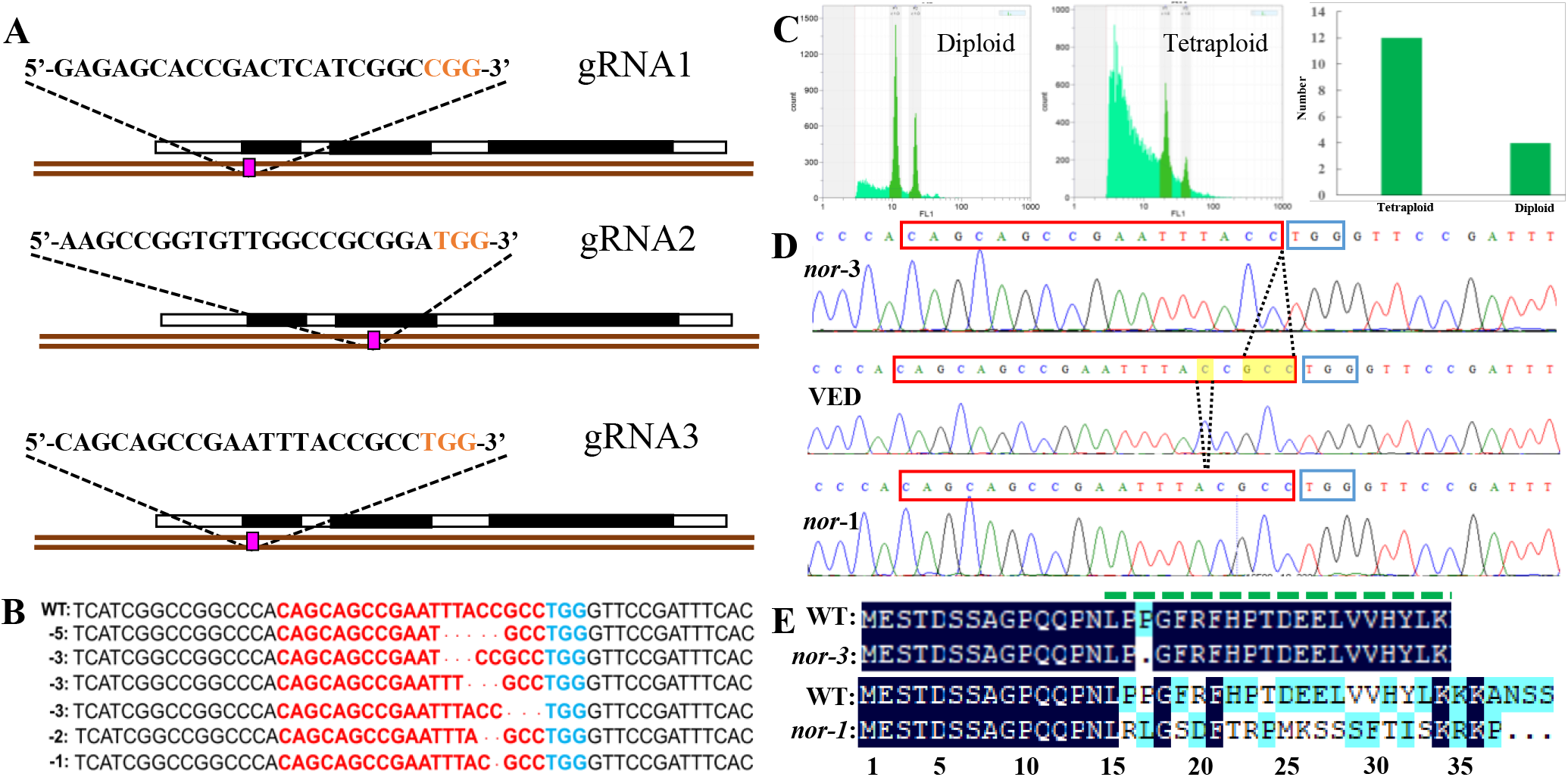
Generation of *CmNAC-NOR* disrupted mutant lines by CRISPR/Cas9. **A** Illustration of the *CmNAC-NOR* gene structure and the three gRNAs in exons 1 and 3. **B** Six independent mutations at gRNA3 targeting sites obtained. **C** Ploidy test of transgenic lines. Left, diploid line NOR-g3-22; Middle, tetraploid line NOR-g3-20; Right, total number of diploid and tetraploid plants obtained. **D** Validated edited lines by Sanger sequencing. The yellow boxes and dotted lines indicate the deleted nucleotides and their position. **E** Amino acid alignment of the CmNAC-NOR gRNA3-target region in VED (WT), *nor-3* and *nor-1*. NAC subdomain A is shown by a dashed green line above the protein sequences.

### Ethylene production and volatile profile in *nor* mutants

Given the key role of ethylene on initiating climacteric fruit ripening (McMurchie et al., 1972), we first compared the ethylene production between controls and both *nor* mutants. The ethylene production was recorded starting from 25 days after pollination (DAP), and the results showed a significant delay (~8 days) in the production of ethylene in *nor-3* when compared to VED and NE, but without a significant difference in the amount of ethylene produced (Figure 2A). For *nor-1* mutant in homozygosis, we did not detect any ethylene production even at 65 DAP (Figure 2A). Interestingly, the *nor-1* allele in heterozygosis showed a ~20 days delay in ethylene production, and the amount of ethylene produced was much lower than the controls (Figure 2A).

**Figure 2.**
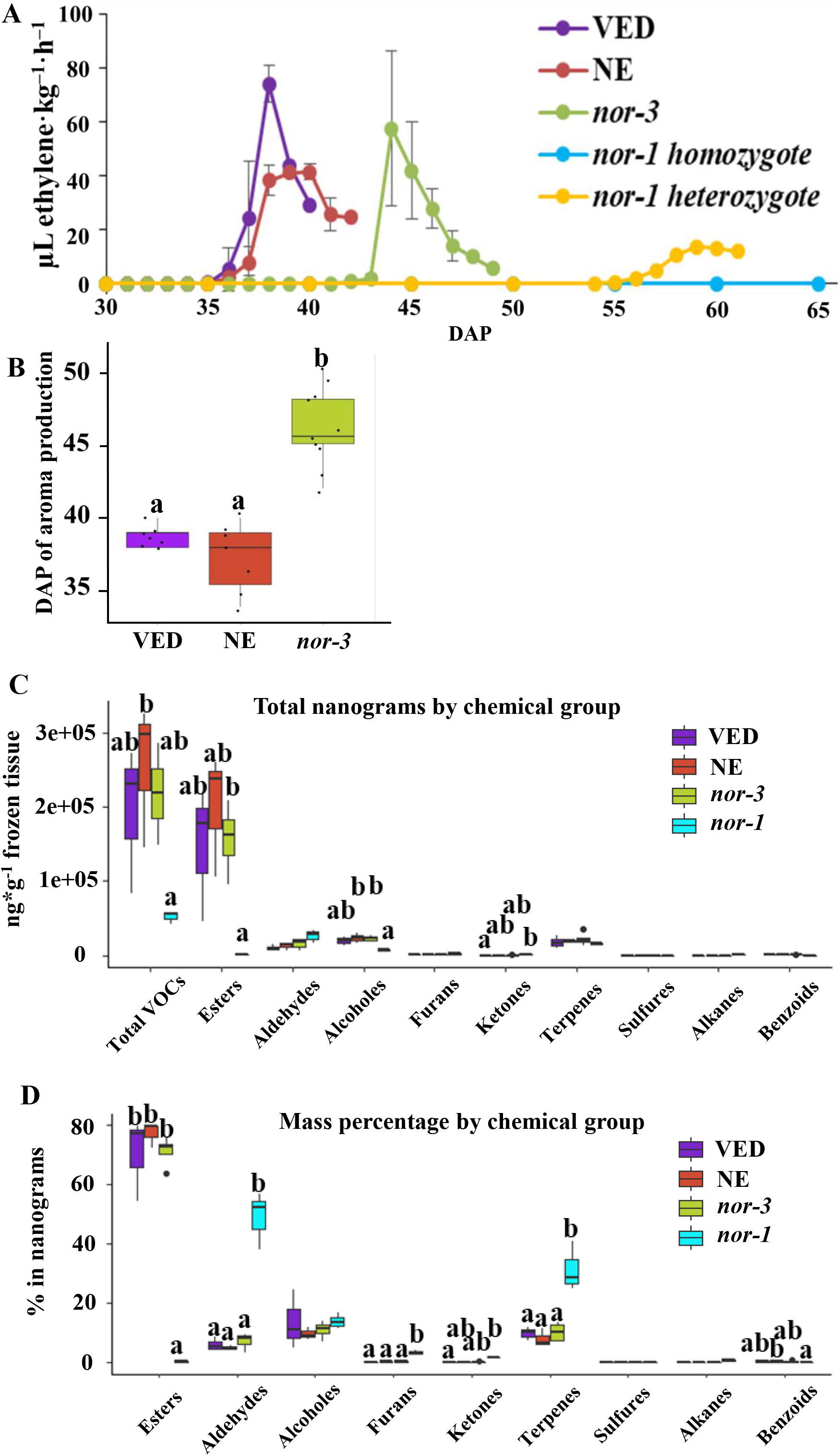
Ethylene production and volatile profile in *nor* mutants. **A** Ethylene production in VED, NE (non-edited control), *nor-3, nor-1 homozygote*, and *nor-1 heterozygote* according to days after pollination (DAP). Means are plotted ±SD (n=5) except *nor-1* heterozygote (n=1). **B** Phenotypic comparison of aroma production in VED, NE, and *nor-3* at harvest. Lower case letters indicate significant differences (P<0.05, n>5). **C-D** Volatile profile in VED, NE, *nor-3*, and *nor-1* at harvest estimated by mass (total nanograms, C) and mass percentage (D). Lower case letters indicate significant differences (P<0.05, n=3).

VED is a *cantalupensis* melon, which produces intense aroma during climacteric fruit ripening due to the high production of esters (Mayobre et al., 2021; Obando-Ulloa et al., 2008). Therefore, we also compared the aroma production between controls and both *nor* mutants. In *nor-3* fruits, the aroma production was significantly delayed compared to the controls at the ripe stage (Figure 2B). However, we couldn’t detect aroma production in *nor-1*. To investigate which volatiles were altered in both mutants, we performed GC-MS to study the fruit flesh volatile profile. As shown in Figure 2C-D, we did not find significant differences in the volatile profile between *nor-3* and controls, but *nor-1* mutants showed a completely different profile with an important decrease in total VOCs produced, which explains the lack of smell by olfactory evaluation. In *nor-1* mutant, we detected much less ester compounds and an increase in aldehydes, furans and terpenes compared to the wild type VED (Supplementary Table S1).

### Partially disrupting *CmNAC-NOR* in *nor-3* delays fruit ripening but does not affect fruit quality

The results from ethylene and aroma production suggest that *nor-3* has a delayed ripening phenotype, and similar results were obtained with other ripening related traits. Flesh color of *nor-3* at 40 and 49 DAP is similar to that of NE at 32 and 40 DAP, respectively (Figure 3A), confirming the 8-9 days ripening delay in *nor-3*. In addition, days of abscission layer formation (Figure 3B) and rind color change (Figure 3C) of *nor-3* fruit were also significantly delayed than those of controls at the ripe stage. However, we could not see a significant difference in the amount of ethylene production (Figure 2A), which is consistent with our previous findings in *CmNAC-NOR* TILLING mutants (Ríos et al., 2017). We also measured soluble solids content (SSC, Figure 3D), fruit weight (Figure 3E) and flesh firmness (Figure 3F) at harvest, and no significant differences were detected among NE, *nor-3* and VED.

**Figure 3.**
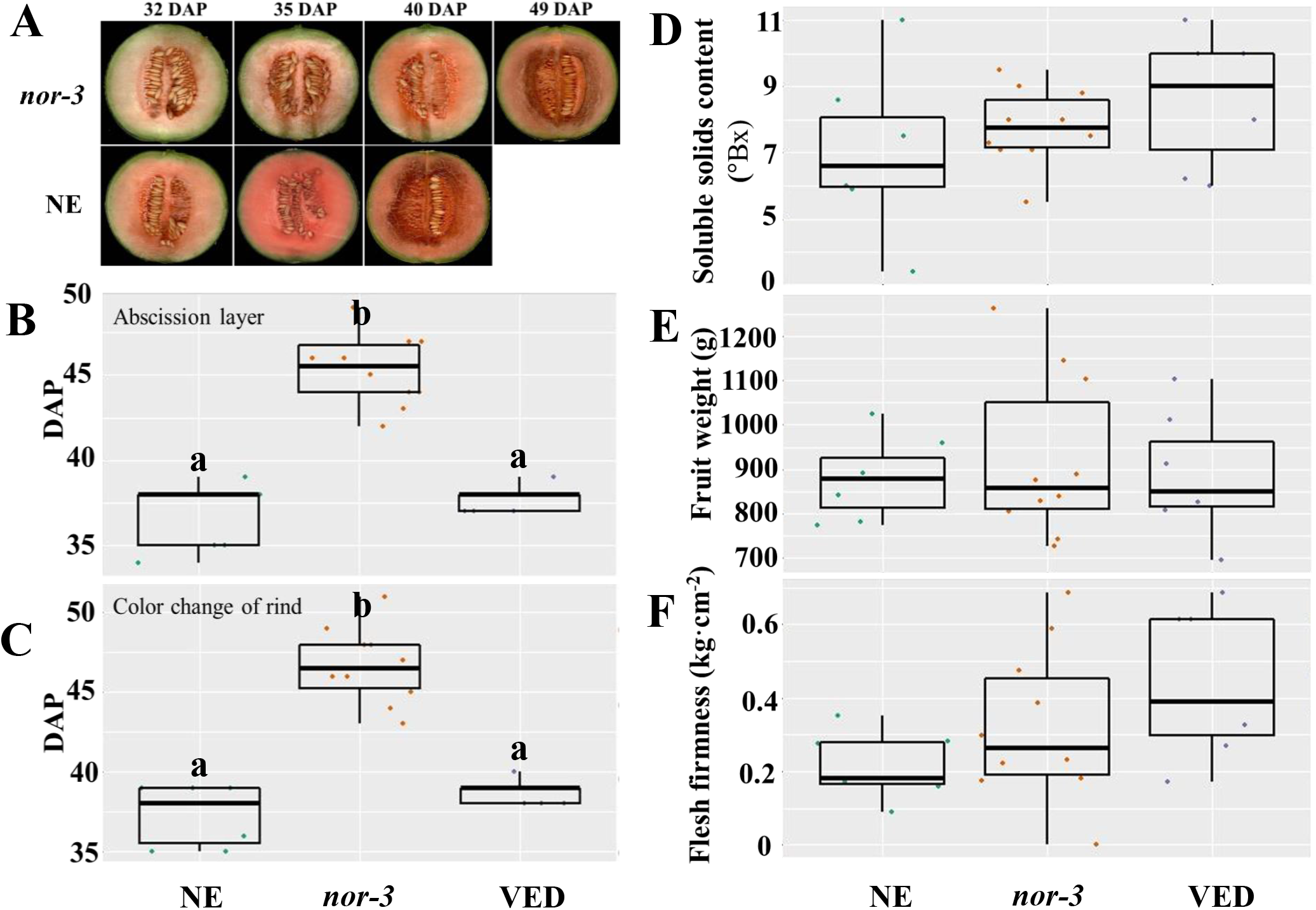
Partially disrupting *CmNAC-NOR* in *nor-3* delays fruit ripening. **A** Fruit ripening phenotype of *nor-3* and NE (non-edited control) under natural ripening conditions at different days after pollination (DAP). **B-F** Phenotypic comparisons according to abscission layer formation (**B**), color change of the rind (**C**), soluble solids content (**D**), fruit weight (**E**) and flesh firmness (**F**) among NE, *nor-3* and VED. Lower case letters indicate significant difference (P>0.05). Means are plotted ±SD (n>5).

### *CmNAC-NOR* knock-out in *nor-1* blocks climacteric ripening

The *nor-1* mutant showed a different behavior in fruit ripening than the *nor*-3 mutant, as the progress of climacteric ripening was totally blocked in *nor-1*, in contrast to only being delayed in *nor-3*. Ethylene production, rind color change, aroma production and abscission layer formation did not occur in *nor-1* (Figure 4A). As shown in Figure 4B, the external color of VED changed to yellow at 38 DAP, while the rind color of *nor-1* remained green at 78 DAP. We also compared flesh color and carotenoid content. *nor-1* flesh was slightly less orange than VED (Figure 4B), however the total carotenoid content was nor significantly different (data not shown). Surprisingly, *nor-1* seeds were not well developed (Figure 4C), resulting in an extremely low germination rate (1.25 %) (Figure 4D and E). However, seed development was not affected in *nor-3* (Figure 4C), which had an 82.86% germination rate (Figure 4D and E). In addition, flesh firmness of *nor-1* was harder than that of VED and NE fruits (Figure 4F). We could not see a significant difference in SSC (Figure 4G) and fruit weight (Figure 4H) among VED, NE and *nor-1* fruits.

**Figure 4.**
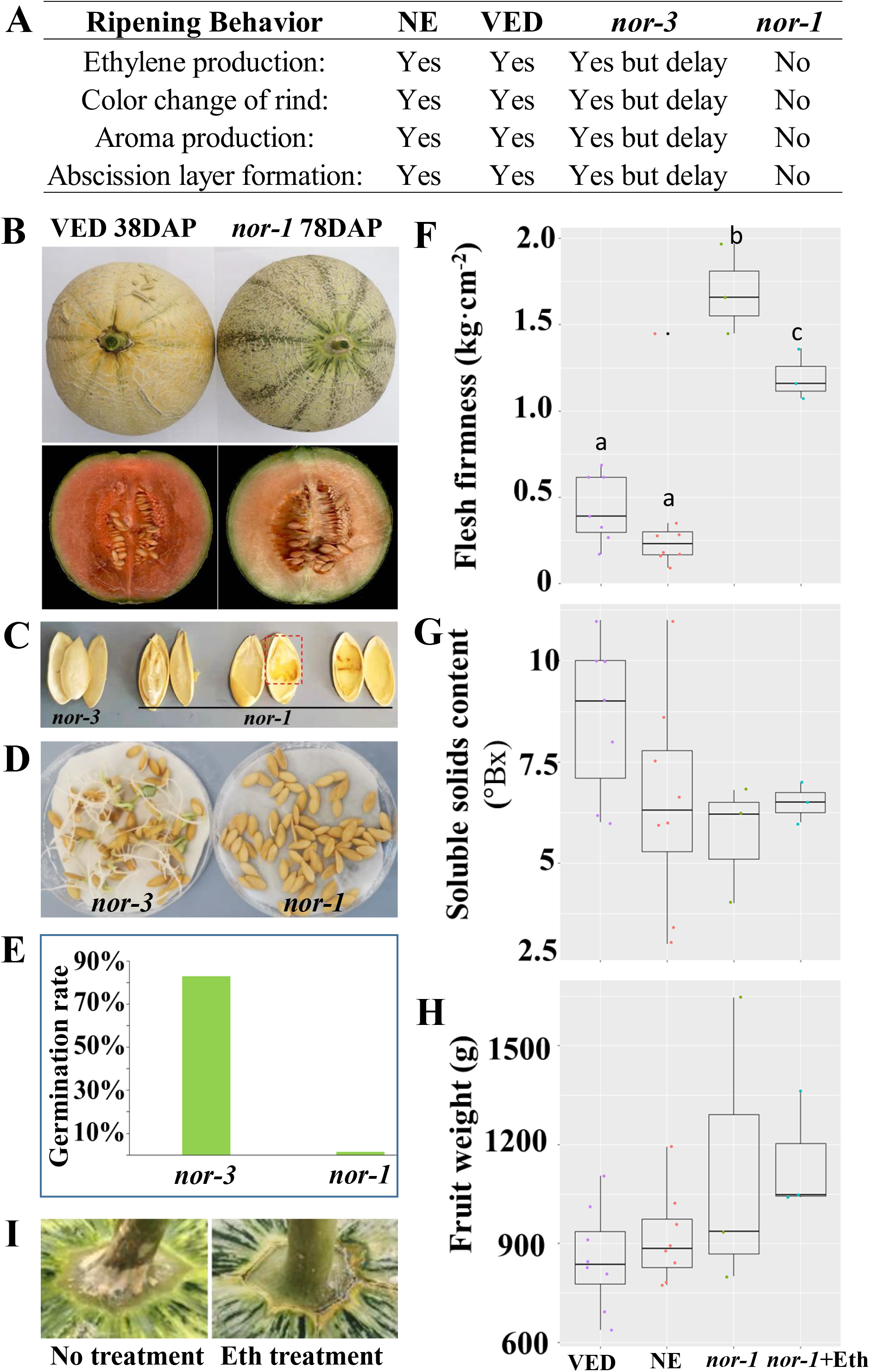
*CmNAC-NOR* knock out in *nor-1* blocks climacteric ripening. **A** Difference of ripening behavior among NE, VED, *nor-3* and *nor-1*. **B** Fruit rind and flesh of VED and *nor-1* under natural ripening conditions. **C** Phenotypic comparison of seeds between *nor-3* and *nor-1* at 38 and 78 DAP respectively. **D-E** Germination efficiency of *nor-3* and *nor-1* seeds. **F-H** Phenotypic comparison according to flesh firmness (**F**), soluble solids content (**G**) and fruit weight (**H**) among NE, VED and *nor-1. nor-1*+Eth: *nor-1* fruit after ethylene treatment. **I** Abscission layer of *nor-1* appears after ethylene treatment.

### Ethylene treatment did not recover climacteric ripening in *nor-1*

Given that ethylene plays a major role in the ripening of climacteric fruit (Ayub et al., 1996), we explored if an external ethylene treatment could induce ripening in *nor-1* mutants. The results showed that climacteric ripening could not be induced after one-week of external ethylene treatment. Rind color did not change to yellow (Figure 4B), fruits did not produce aroma, and flesh firmness slightly decreased but was still significantly harder than VED an NE fruits (Figure 4F). We did not see a significant change in SSC (Figure 4G) and fruit weight (Figure 4H). However, the abscission layer was induced at 3 days after ethylene treatment (Figure 4I).

## Discussion

NAC transcription factors are a large gene family involved in plant development and environment stress response (Guo and Gan, 2006; Hernández and Sanan -Mishra, 2017; Nuruzzaman et al., 2013). Among them, the *SlNAC-NOR* gene from tomato is well known as a key regulator in fruit ripening (Giovannoni, 2007). In previous studies we characterized *CmNAC-NOR* as a homologue of *SlNAC-NOR*, and proved its involvement in fruit ripening with the delayed ripening phenotype of two TILLING mutant lines (Ríos et al., 2017). In this study, we have obtained two diploid *CmNAC-NOR*-disrupted melon plants with different edited sites (*nor-1* and *nor-3*) by using the CRISPR/Cas9 system (Figure 1). *nor-1* is a complete knock-out mutant whose climacteric ripening behavior is almost absent, without ethylene and aroma production, a volatile profile similar to unripe fruit, absence of abscission layer, and no external color change (Figures 2 and 4), suggesting that *CmNAC-NOR* plays a significant role on regulating fruit ripening in melon. The *nor-3* is a knock-down mutant with one amino acid deletion at the NAC (<u>N</u>AM, <u>A</u>TAF1,2, <u>C</u>UC2) domain, which causes around 8 days delay in ripening, but without affecting fruit quality at the ripe stage (Figure 3), suggesting a potential way to control fruit ripening in melon by differently disrupting *CmNAC-NOR*. Moreover, the heterozygous *nor-1* also showed a 20 days delay in ethylene production (Figure 2A), suggesting that *nor-1* might have potential for creating long shelf life fruit in melon breeding programs.

In tomato, fruit ripening was affected in both natural and CRISPR/Cas9 knock out mutants of *SlNAC-NOR* (Gao et al., 2020). However, the fruit of *SlNAC-NOR* mutations (*CR-NOR*) induced by CRISPR/Cas9 showed a much less severe phenotype than the natural mutant *slnac-nor* (Gao et al., 2020; Wang et al., 2019). Mature fruit of *slnac-nor* mutant did not produce an ethylene burst (Adaskaveg et al., 2021; Giovannoni, 2007), and had little carotenoid content (Giovannoni et al., 1995; Kumar et al., 2018), but *CR-NOR* fruits can produce ethylene, and synthesize much more carotenoids than *slnac-nor* at the ripe stage (Gao et al., 2020). Therefore, *slnac-nor* was reported as a gain-of-function mutation (Gao et al., 2020) suggesting that *SlNAC-NOR* did not act as a master regulator but as a major gene controlling the ripening process (Wang et al., 2020a). In climacteric melon such as VED, the fruit ripening occurred around 35~40 DAP (Fig. 2A), and was associated with a transient increase in autocatalytic ethylene production, accompanied by changes in rind and flesh color, flesh firmness, sugar content and aroma production (Mayobre et al., 2021; Pereira et al., 2018; Pereira et al., 2020). In the complete knock-out mutant *nor-1*, ethylene production was blocked (Fig. 2A), the aroma component changed (Fig. 2C), the rind color did not change from green to yellow (Figure 4B), no abscission layer occurred, and the flesh was harder than VED (Figure 4F). The flesh color of *nor-1* seemed visually less orange than VED (Figure 4B), although we did not detect significant differences in carotenoid content. Our findings suggest that *nor-1* resembles both the tomato phenotypes of *slnac-nor* and *CR-NOR* mutants. Unlike the tomato CRISPR mutant *CR-NOR*, the main climacteric ripening components were almost blocked in *nor-1* melon. In addition, *nor-1* was insensitive to external ethylene treatment (Figure 4), except for abscission layer formation, suggesting that *CmNAC-NOR* gene is a major key regulator of fruit ripening in melon.

The different phenotypes observed between tomato and melon CRISPR *NAC-NOR* mutants might be explained by their different editing patterns. Although all of them lost the transcriptional regulation region, their NAC domain is altered at different levels. The NAC domain contains five subdomains (A-E) that play an important role in DNA-binding (Ernst et al., 2004; Kikuchi et al., 2000). The *slnac-nor* mutant contained a complete NAC domain, resulting in a gain-of-function mutation; the *CR-NOR* mutant produced a truncated protein of 47 aa, which lost NAC subdomains B-E, but still had NAC subdomain A (Gao et al., 2020); in our work, the editing of *nor-1* started from NAC subdomain A (Figure 1E), thus the whole NAC domain was affected in *nor-1*, resulting in a loss-of-function mutation.

Fruit flavor is an important trait as it affects consumer habit. Volatile esters are major contributors to fruit flavor giving the fruity aroma to climacteric melons (El Hadi et al., 2013). Compared to the controls, the esters content was dramatically reduced in *nor-1* and the content of aldehydes was increased, which explain the green, fresh aroma of these fruits. *nor1* VOCs profile was more similar to unripe melons or to non-climacteric melons such as *inodorus* types (Mayobre et al., 2021) than to the VED profile. These results suggest that *CmNAC-NOR* could be involved in the regulation of the *AAT* genes, which are known to be ethylene-dependent and are responsible for the volatile ester formation (Cao et al., 2021; El-Sharkawy et al., 2005). This is also consistent with a recent study that reported that NAC transcription factor *PpNAC1* (with homology to *SlNAC-NOR*) regulates fruit flavor ester biosynthesis in peach by activating ripening-related *AAT* expression (Cao et al., 2021).

An unexpected phenotype of *nor-1* was that seeds were not well developed (Figure 4C). This phenotype was not previously reported in the tomato *NOR* mutant or in other species, such as peach (Pirona et al., 2013), apple (Yeats et al., 2019), and strawberry (Martín -Pizarro et al., 2021). However, there are some reports suggesting that NAC transcription factors regulate seed development and play a role in seed germination (Kim et al., 2008; Park et al., 2011; Wang et al., 2021). In a recent study, knock out of *ClNAC68* gene in watermelon delayed seed maturation and germination, but the germination rate was not affected (Wang et al., 2021), suggesting that there are additional NAC genes with diverse functions that regulate seed development.

In this study, we provided evidence that supports *CmNAC-NOR* as a key player in regulating climacteric fruit ripening in melon. As a master regulator, *CmNAC-NOR* independently mediates many ripening associated traits. Our findings also suggest that *CmNAC-NOR* can be a potential target in breeding programs to modulate fruit maturation and shelf life in melon.

## Materials and methods

### Plant material and generation of constructs

The cantaloupe inbred line VED (climacteric) was used in this study. For editing *CmNAC-NOR* in VED, three gRNAs (gRNA1, gRNA2, and gRNA3; Supplementary Table 2) were designed based on the genomic sequence of *CmNAC-NOR* using Breaking-Cas (Oliveros et al., 2016). Afterwards, gRNA1 and gRNA2 were inserted into the vector pBS_KS_Bsa_Bbs_tandem with the *BbsI* and *BsaI* sites, respectively, then cut by restriction enzymes *SpeI* and *KpnI*, and inserted into the final vector pB7-CAS9-TPC to obtain the gRNA1-gRNA2-CAS9 construct. The same protocol was used to generate the gRNA2-gRNA1-CAS9 and gRNA3-gRNA1-CAS9 constructs. Afterwards, the constructs were transformed into Agrobacterium (Agl-0) and identified by cloning PCR. Cloning vectors were kindly provided by Prof. Puchta (KIT, Germany).

### Melon transformation

Melon transformation was performed by a cotyledon transformation method as described by Castelblanque et al, 2008 with modifications as in (García -Almodóvar et al., 2017). In brief, half of the proximal parts of the cotyledons from 1-day-old seeds were cut and co-cultured with transformed Agrobacterium. The inoculated explants were placed on regeneration media and then in selection media in the presence of DL-Phosphinothricin (PPT). Every 3-4 weeks, the green cluster buds were cut and explants were moved to fresh selection media. When the regenerated shoots grew high, they were cut, separated from the explants, and put into big test tubes individually in rooting media. When the rooted plantlets were big enough, a leaf section was cut to identify edited T0 plants.

### Genotyping and ploidy test

Genomic DNA was extracted from young leaves of melon plants by an improved CTAB method (Pereira et al., 2018). To genotype the candidate plants, *CAS9* gene was amplified to confirm that plants were transgenic, then the target region of the three gRNAs was amplified and sequenced. Primers used in this study are listed in Supplementary Table 2. At the same time, young leaves were harvested and sent to Iribov (Heerhugowaard, Netherlands) to do the ploidy test by flow cytometry (FCM).

### Identification of CAS9 free T_1_ plants

Diploid T_0_ plants that carried the *CAS9* gene were selected, grown and self-pollinated to obtain the T_1_ seeds. Afterwards, T_1_ seeds were germinated and genotyped, and the *CAS9* free plants with or without *CmNAC-NOR* editions were selected for further experiments.

### Fruit phenotyping

Fruit quality traits, especially those associated to climacteric ripening behavior were assessed after harvest as previously described (Pereira et al., 2018; Ríos et al., 2017). In brief, the production of ethylene in the fruits was measured from 25 days after pollination (DAP) by using a non-invasive ethylene quantification method (Pereira et al., 2017), until fruit dropped or 65 DAP when it did not drop. The same sealing method for measuring ethylene (Pereira et al., 2017) was used for ethylene treatment, where250 ml of 50 p.p.m ethylene was injected in the bag with a syringe, keep close for one week, and then phenotyping was performed after opening the bag. The production of aroma was detected by olfactory evaluation of fruits from 25 DAP until harvest. The days for abscission layer formation were also recorded. External color change during fruit ripening was phenotyped visually. Fruits were weighed at harvest. Soluble solid content was analyzed with a digital hand refractometer (Atago Co. Ltd., Tokyo, Japan). Flesh firmness was measured using a penetrometer (Fruit TestTM, Wagner Instruments).

### Volatiles analysis

Two grams of frozen grinded flesh were weighed and added to 20 ml chromatography vials with 1 g of NaCl and 7 ml of saturated NaCl solution containing 15 ppm of 3-hexanone as internal standard. Samples were stored at 4 ºC a maximum of 7 days. Solid - Phase Micro-Extraction (SPME) was carried out by pre-heating samples for 15 min at 50 ºC and 250 rpm. The SPME fiber (50/30 µm DVB/CAR/PDMS, Merck®, Darmstadt, Germany) was exposed to the vial headspace for 30 min. Splitless injection was performed into a 7890A gas chromatograph (GC) equipped with a Sapiens-X5MS capillary column (30 m/0.25 mm/0.25 µm, Teknokroma®, Sant Cugat del Vallès, Spain), with 10 min of thermal desorption at 250 ºC. Oven was set to 50 ºC for 1 min, then increasing 5 ºC/min to 280 ºC and holding 5 min. Carrier gas was helium at head pressure of 13.37 psi. A mass spectrometer (MS) 5975 C (Agilent Technologies®, Santa Clara, CA, U.S.) was coupled to the GC, with a source temperature of 230 ºC and quadrupole temperature was set to 150 ºC. We performed an untargeted analysis and volatiles were identified by comparison of their mass spectra with the NIST 11 library (NIST/EPA/NIH) and by their Kovats retention index, calculated using a mix of alkanes (C7-C40 in hexane, Merck®, Darmstadt, Germany) under the same chromatographic conditions. The content of each volatile was calculated comparing the peak area to the internal standard peak. A Shapiro-Wilk normality test was conducted (α = 0.05) and a multiple variable t-test was performed. Wilcoxon test was performed to compare edited plants against wild type.

### Data analysis

DNA and protein sequence alignments were obtained with DNAMAN version 7. All the statistical analyses were obtained using the software R (v3.5.3) (https://www.r-project.org/). ANOVAs and pairwise t-test were performed using R package “rstats”. In general, significance was fixed at p-value <0.05.

## Acknowledgements

This work was supported by the Spanish Ministry of Economy and Competitiveness grants AGL2015–64625-C2–1-R and RTI2018-097665-B-C2, the Severo Ochoa Programme for Centres of Excellence in R&D 2016-2010 (SEV-2015-0533), the CERCA Programme/Generalitat de Catalunya and 2017 SGR 1319 grant from the Generalitat de Catalunya to JGM. LB was also supported by grants from Youth Project of National Natural Science Foundation of China (31902035), The International Postdoctoral Exchange Fellowship Program of China (20170053) and a postdoctoral grant from the Severo Ochoa Programme for Centres of Excellence in R&D 2016-2010 (SEV-2015-0533). MSD was supported by a grant from the Spanish Ministry of Economy and Competitiveness. CM was supported by a grant from the Secretaria d’Universitats i Recerca del Departament d’Empresa i Coneixement de la Generalitat de Catalunya and the co-funding of the European Social Fund (ESF-“ESF is investing in your future”. Special thanks to Fuensanta García and Elena del Blanco for technical assistance in field and lab operations and to Laura Valverde for carotenoid measurements.

## Author contributions

LB performed the experimental work, data analysis and obtained the original draft of the manuscript. MSD grew the plants and phenotyped the ripening behavior and ethylene measurements. CM performed the volatile experiments. AMM-H supervised the CRISPR-Cas9 and genetic transformation experiments. MP and JG-M designed and supervised the work and reviewed and edited the manuscript.

## Supporting Information

**Supplementary Table 1**. VOCs detected in melon flesh samples.

**Supplementary Table 2**. List of primers and their uses.

